# Bi-level positive airway pressure (biPAP) for non-invasive respiratory support of foals

**DOI:** 10.1101/2020.11.10.376392

**Authors:** Sharanne L Raidal, Lexi Burgmeestre, Chee Sum Melanie Catanchin, Chris T Quinn

## Abstract

Respiratory insufficiency and pulmonary health are important considerations in equine neonatal care, as the majority of foals are bred for athletic function. The administration of supplementary oxygen is readily implemented in equine practice settings, but this does not address respiratory insufficiency due to inadequate ventilation and is no longer considered optimal care for hypoxia in some settings. Non-invasive ventilatory strategies including continuous or bi-level positive airway pressure are effective in human and veterinary studies, and may offer improved respiratory support in equine clinical practice. The current study was conducted in two parts to investigate the use of a commercial bilevel positive airway pressure (biPAP) ventilator, designed for home care of people with obstructive respiratory conditions, for respiratory support of foals. In Part 1 a prospective observational study was conducted to evaluate the effect of sequential application of supplementary oxygen and then biPAP for respiratory support of five foals ≤ 4 days of age hospitalised with respiratory in sufficiency (Group 1) and four healthy, sedated foals < 28 days of age (Group 2). In Part 2, biPAP and supplementary oxygen were administered to six healthy foals with pharmacologically induced respiratory insufficiency in a two sequence, two phase, cross-over study (Group 3). Non-invasive ventilation by biPAP improved gas exchange and mechanics of breathing (increased tidal volume, decreased respiratory rate and increased peak inspiratory flow) in foals, but modest hypercapnia was observed in healthy, sedated foals (Groups 2 and 3). Clinical cases (Group 1) appeared less likely to develop hypercapnia in response to treatment, however the response in individual foals was variable, and close monitoring is necessary. Clinical observations, pulse oximetry and CO_2_ monitoring of expired gases were of limited benefit in identification of foals responding inappropriately to biPAP, and improved methods to assess and monitor respiratory function are required in foals.

## Introduction

Respiratory disease has long been recognised as of considerable economic importance in newborn foals (1), and as an important cause of morbidity and death in neonates presented for veterinary care (2, 3). Optimal respiratory support is highly desirable to optimise survival and preserve respiratory function in animals bred largely for their athletic potential.

The use of non-invasive ventilation (NIV) is now widely regarded as the most effective approach for respiratory support of human neonates (4, 5), with continuous positive airway pressure (CPAP) shown to reduce the number of preterm infants requiring admission to neonatal intensive care (6), and to decrease the risk of bronchopulmonary dysplasia or death in neonates requiring respiratory support (7). The technique involves the delivery of a constant positive (greater than atmospheric) pressure to the airway and preserves spontaneous respiration. The physiological effects are complex and likely to vary depending on the underlying pathology (8), but benefits have been attributed to increased functional residual capacity, decreased work of breathing and reduced airway resistance (4). Previous studies have demonstrated that CPAP is associated with improved respiratory function in a number of veterinary species (9–12). CPAP has recently been shown to improve gas exchange in healthy foals with pharmacologically induced respiratory suppression (13), however hypercapnia was observed in treated foals in this study, and has been observed previously in anaesthetised horses during CPAP (10, 11, 14).

Bi-level positive airway pressure (biPAP) is also recognised for the management of respiratory insufficiency in human neonates, and has recently demonstrated improved treatment outcomes in preterm human neonates in comparison to CPAP (15, 16). By using lower expiratory pressures, biPAP promises improved expiratory function and is recommended for management of conditions associated with hypercapnia, such as chronic obstructive airway disease or asthma (17–19). In human patients with obstructive airway conditions, expiratory airflow limitations may cause increased PaCO_2_ due to overdistension of alveoli and consequent increased alveolar dead space (20–23), an effect which has been termed dynamic hyperinflation (24).

Whereas human neonates exhale passively (25), both inspiration and expiration are active processes, requiring muscular effort in foals (26). Equids might therefore be predisposed to expiratory flow limitations and retention of CO_2_ if active breathing strategies are unable to overcome expiratory pressures during CPAP.

We hypothesised that CPAP might be associated with diffuse expiratory flow limitation, increased intrinsic positive end-expiratory pressure (PEEP*i*) and alveolar overdistension in foals, and this would then predispose to hypercapnia. Lower expiratory pressures during biPAP might then be expected to facilitate expiration and ameliorate this effect. The current study was undertaken to determine whether a commercially available bi-level respiratory device available for the home care of people with respiratory disease might offer improved ventilatory support in healthy foals with pharmacologically induced respiratory disease, and to characterise the response of hospitalised neonatal foals with respiratory insufficiency managed with biPAP. We hypothesised that biPAP would be associated with benefits in improved gas exchange previously observed during CPAP (Raidal et al), but with less CO_2_ retention. Intentionally, the study was designed to evaluate low cost intervention and monitoring strategies that might be safely implemented in equine practice or on farm.

## Materials and Methods

### Animals

The study was conducted in two parts: an observational study of foals < 4 weeks of age (Groups 1 and 2), and an interventional study (Group 3). The study protocol was approved by the Charles Sturt University Animal Care and Ethics Committee (ACEC A18044).

Group 1 consisted of five foals (Table 1) presented to the equine neonatal intensive care unit (NICU) at the Veterinary Clinical Centre, Charles Sturt University, with respiratory insufficiency defined as hypoxia (PaO_2_ <85 mmHg) and/or hypercapnia (PaCO_2_ > 50 mmHg). Four had clinical signs consistent with neonatal multisystemic maladjustment syndrome (NMMS) including recumbency (F1, F3, F4), altered mentation (F1, F3, F4, F5), and failure to nurse (F1, F3, F4, F5). One university owned foal (F2) was presented with respiratory insufficiency and polysynovitis (all four fetlock joints, both inter-carpal joints and both subcutaneous calcaneal bursae) at 12 hours post-partum. Haematology and serum biochemistry were unremarkable for this foal, and synovial fluid was cytologically normal. A second university-owned foal (F5), born after 406 days gestation, with characteristics of intra-uterine growth retardation and dysmaturity (small size, dished face, floppy ears, flexor laxity) required assistance to nurse. She had episcleral haemorrhage at birth, and synovial effusion of both carpal sheaths within 12 h. Haematology and serum biochemistry were unremarkable, and she responded well to supportive care (intravenous fluids, supplementary feeding via an indwelling nasogastric tube, plasma transfusion, shoe extensions) on farm over 48 hours, but was represented at four days of age having been found submerged in a water trough. Other foals were presented within 48 hours of birth.

**Table 1:**
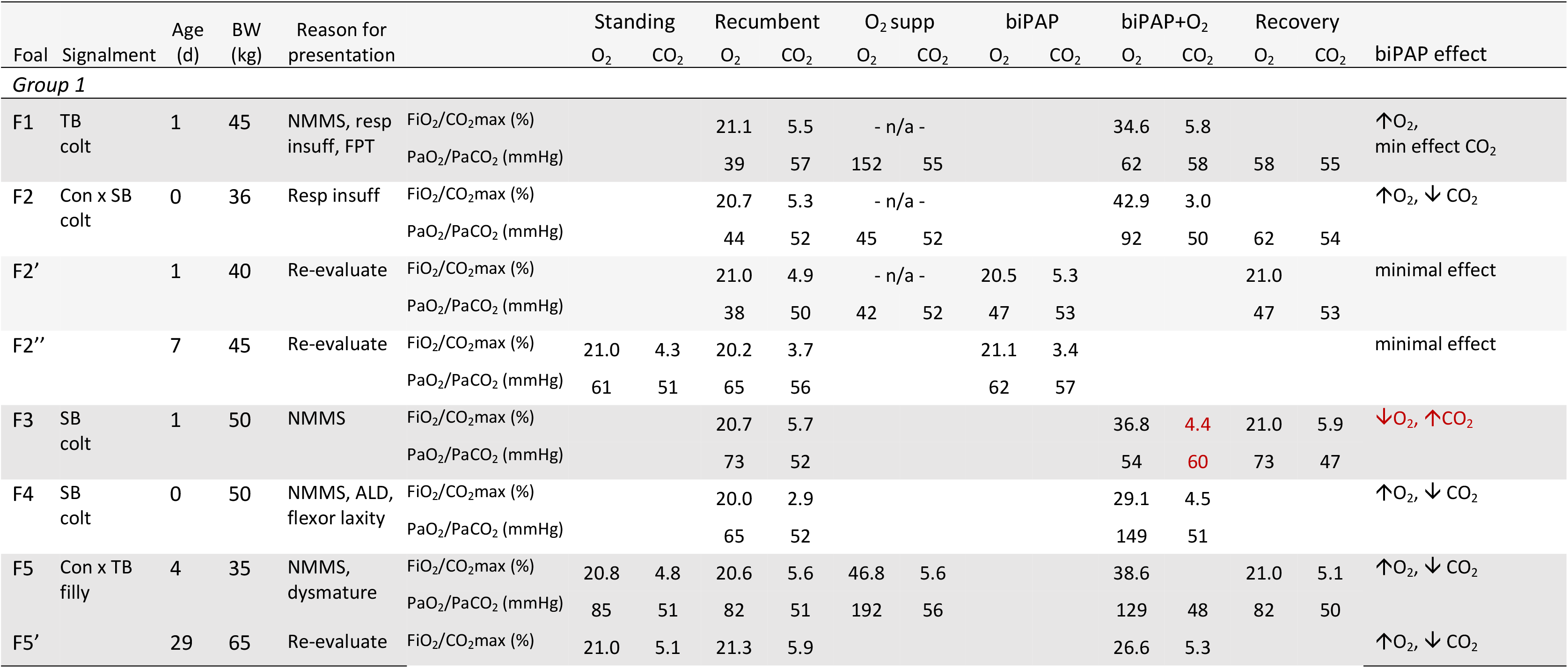
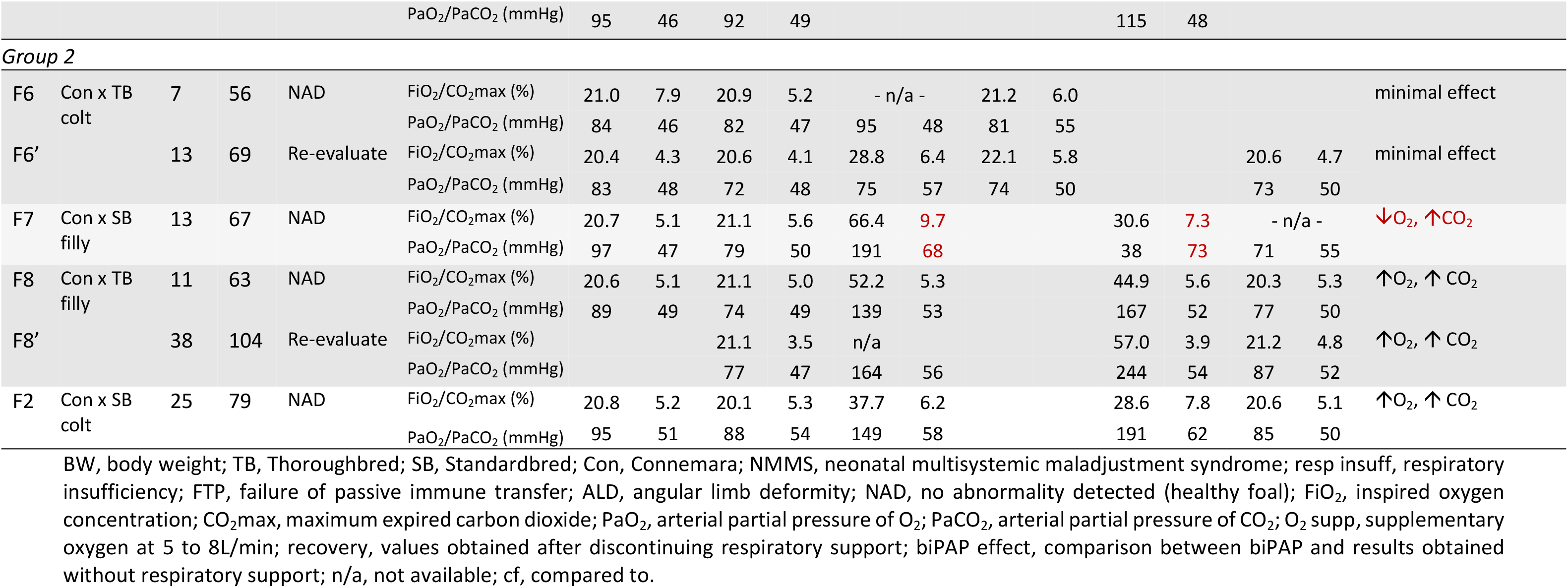
Foals available for inclusion in observational studies assessing non-invasive respiratory support. Group 1 foals were hospitalised for treatment of multiple problems including respiratory insufficiency. Group 2 foals were healthy foals < 28 days of age. A number of foals (F2, F5, F6 and F8) were evaluated on multiple occasions. Arterial blood gas (ABG) samples were obtained from all foals at baseline (ambient conditions, standing and/or recumbent), following respiratory support with oxygen supplementation (O_2_ supp) by nasal insufflation (F1) or mask and bi-level positive airway pressure (biPAP) with (+O_2_) or without oxygen supplementation. For Group 1 foals, oxygen flow was 4 L/min (F4), 5 L/min (F5, F5’), 6 L/min (F3) or 7 L/min (F1, F2, F2’) and biPAP settings were 4/20 cm H_2_O (expiratory / inspiratory pressure: F1, F2, F2’), 5/15 cm H_2_O (F2’’, F3), 4/15 H_2_O (F4, F5, F5’). For Group 2 foals, oxygen flow was 8 L/min for F8’, or 5 L/min for all other foals; biPAP pressures were 5/15 H_2_O for all foals except F5’ (4/15 cm H_2_O). Group 1 foals were manually restrained in lateral recumbency; Group 2 foals were sedated prior to restraint in lateral recumbency.

Group 2 foals consisted of four university owned foals aged between 7 and 25 days on initial assessment, with no abnormalities on veterinary examination (Table 1).

Group 3 comprised six research foals (two colts, four fillies) of mean age 47.7 days (range 44 to 52 days) and mean body weight 111.2 kg (range 86 – 125 kg). All foals were Connemara cross breeding. Four were born unassisted with no abnormalities during gestation or parturition. Two foals (F1 and F6) were included after responding positively to supportive care as described above. All Group 3 foals were normal on veterinary examination at the time of recruitment into the study, and prior to each intervention.

### Group 1 (foals hospitalised for treatment of respiratory insufficiency)

Group 1 foals were included with owner consent if they had evidence of respiratory insufficiency defined as hypoxaemia (PaO_2_ < 85 mmHg) or hypercapnia (PaCO_2_ > 50 mmHg). Blood gas analysis (GEM Premier, Model 3500; Abacus ALS, Macquarie Park, Australia) was performed on anaerobically collected samples to determine partial pressures of oxygen and carbon dioxide (PaO_2_ and PaCO_2_, respectively), haemoglobin saturation (sO_2_) and pH. At the discretion of treating veterinarians, foals were assessed sequentially breathing room air, following administration of supplementary O_2_ (5 L/min) by nasal insufflation or via a large veterinary anaesthesia mask (Size 4, 115 / 43 mm; VetQuip, Eastern Creek, NSW), and following administration of biPAP with or without supplementary O_2_. Respiratory support (biPAP) was delivered via the veterinary anaesthesia mask connected to a vented non-rebreathing elbow valve (Oracle 2 Vented Non-rebreathing Valve, 400HC206, Fisher and Paykel Healthcare, Nunawading, Victoria, Australia) and hence to a commercial, bi-level pressure support ventilator specifically designed for non-invasive mask ventilation (VPAPTM III ST, ResMed Ltd, Bella Vista, NSW) via standard air tubing (ResMed Ltd, Bella Vista, NSW) of two metre length (Supplementary Figure S1). Based on previous findings, a minimum respiratory rate (RR) of 15 bpm was set on the ventilator, such that a breath would be initiated if spontaneous ventilation fell below this rate. Inspiratory pressure (IPAP) was set at 15 cmH2O, and expiratory pressure (EPAP) was set at 5 cmH_2_O. The maximum duration of IPAP was 1.2s to prevent prolonged inspiration, minimum duration of IPAP set at 0.5s to prevent false triggering, and the I:E ratio was 1:2.3. Analysis of inspired and expired gases (FiO_2_ and CO_2_max; PowerLab 4/25, Gas Analyser ML206 and LabChart 8 software; ADInstruments, Bella Vista, NSW) was performed by attaching a gas sampling port to the anaesthetic mask or by inserting a gas sampling tubing into the nares, and following two point calibration with room air (20.9% O_2_, 0.04% CO_2_) and Carbogen (95% O_2_, 5% CO_2_; BOC Gas, Wagga Wagga, NSW). Pulse oximetry (SpO_2_) was performed, when possible, with a transmission probe (Avant 2120, Nonin Medical Inc., Plymouth, MN, USA; distributed by Proact Medical Systems, Port Macquarie, NSW, Australia) placed on the tongue. Results were recorded when the signal was constant over two minutes, pulse rate matched heart rate, and pulse strength was satisfactory. Pulse oximetry was not possible when the anaesthetic mask was in place.

Foals were restrained in lateral recumbency, without sedation, by final year veterinary students on clinical rotation, who were also responsible for holding the mask in place and continuous observation of respiratory rate and effort. Each respiratory intervention was of ≥ 10 minutes duration. Assessments were repeated on two university-owned foals, as shown in Table 1.

### Group 2 (healthy foals)

Group 2 foals were healthy university-owned foals managed in a similar manner to Group 1 foals, assessed sequentially whilst breathing room air, supplementary O_2_ and after administration of biPAP. Assessments were repeated on two foals, as shown in Table 1. Lateral recumbency was induced in Group 2 foals by administration of intravenous injections of diazepam (0.2 mg/kg; Parnell Laboratories, Alexandria, Australia), followed by xylazine (0.02 mg/kg; Illium Veterinary Products, Glendenning, Australia) and fentanyl (5 μg/kg; Hospira, Melbourne, Australia). Each respiratory intervention was of ≥ 10 minutes duration. BiPAP settings were as described for Group 1 foals.

### Group 3 (healthy foals, pharmacological induction of respiratory insufficiency)

A randomised crossover design was used with the first treatment assigned (either biPAP or mask O_2_) determined by coin toss (Supplementary Table S1). Treatment order was reversed for the next data collection day for each foal. The interval between intervention periods ranged from three to six days, and the cross over design included treatment in both left (Phase 1) and right lateral recumbency (Phase 2) for each foal.

Prior to each study, foals were manually restrained for veterinary examination and collection of baseline arterial blood samples from the distal carotid arteries (T-1). A 16G, 89 mm catheter (Terumo Surflo, Macquarie Park, Australia) was placed in the jugular vein of each foal and a baseline sample of venous blood was collected from each foal prior to sedation. Blood gas analysis was performed as described for Group 1 foals. Spirometry was performed on standing foals as previously described (Raidal, McKean et al. 2019) by application of a large veterinary anaesthesia mask (SurgiVet large canine mask, product number 32393B1; Sound Veterinary Equipment, Rowville, Australia) placed on the foal’s muzzle in such a way as to exclude air leaks and to minimise dead space, but not prevent opening of the nares. A respiratory flow head (Respiratory Flow Head 300 L, MLT300L, ADInstruments, Bella Vista, Australia) and gas sampling port were connected to the anaesthesia mask. Dead space of this apparatus was 60 mL (measured by water displacement). Data were collected for up to 60 seconds (sufficient to ensure 10 artefact free breath cycles) in unsedated foals, and analysed using PowerLab 4/25, Gas Analyser ML206 and LabChart 8 software (ADInstruments, Bella Vista, Australia). Tidal volume (Vt), peak inspiratory and peak expiratory air flow (PIF, PEF), and the duration of inspiratory (Ti) and expiratory (Te) phases were determined by post-sampling analysis of six consecutive and artefact free breath cycles representative of tidal breathing. Spirometry, inspired and expired gas analysis (FiO_2_, FeO_2_, FiCO_2_ and FeCO_2_) were performed following calibration of the spirometer pod using a using a seven litre certified calibration syringe (Hans Rudolph Incorporated, Shawnee, Kansas, USA) and the gas analyser was calibrated using a two point calibration of room air (20.9% O_2_, 0.04% CO_2_) and Carbogen (95% O_2_, 5% CO_2_; BOC Gas, Wagga Wagga, Australia). Pulse oximetry (SpO_2_) was performed, when possible, as described for Group 1 foals.

Diazepam (0.2 mg/kg) was administered via the intravenous catheter, and spirometry (T0) was repeated five minutes following treatment. Fentanyl (5 μg/kg; Hospira, Melbourne, Australia) and xylazine (0.02mg/kg; Illium Veterinary Products, Glendenning, Australia) were administered via the jugular catheter, and foals were placed in lateral recumbency. A 22G, 2.5cm polyurethane catheter (Surflo, Terumo Australia Pty Ltd, Macquarie Park, Australia) was placed aseptically into the lateral metatarsal artery, and an arterial blood sample collected anaerobically (T0). Foals were monitored by determination of cardinal signs (HR, RR, temperature, MAP), arterial blood gases (ΡaO_2_, ΡaCO_2_, sO_2_, pH), spirometry and inspired/expired gas analysis. Spirometry data were collected over 20 to 40 s, after collection of arterial blood samples, to minimise any effects attributable to apparatus dead space. All foals were sampled 10 min following administration of diazepam (T0), and respiratory suppression was induced by continuous infusion of fentanyl (0.005 mg/kg/hr) and xylazine (0.7 mg/kg/hr) in 0.9% sodium chloride delivered via syringe pump (Alaris IMED Gemini PC-1 infusion pump; VetQuip Pty Ltd, Erskine Park, NSW), commencing at T0. Samples were again collected after 10 minutes spontaneous respiration (10 minutes following commencement of the fentanyl-xylazine CRI, T1), and respiratory support (biPAP or O_2_ supplementation) was commenced 10 minutes following at this time. The treatment and sample schedule is shown in Table 2.

**Table 2:**
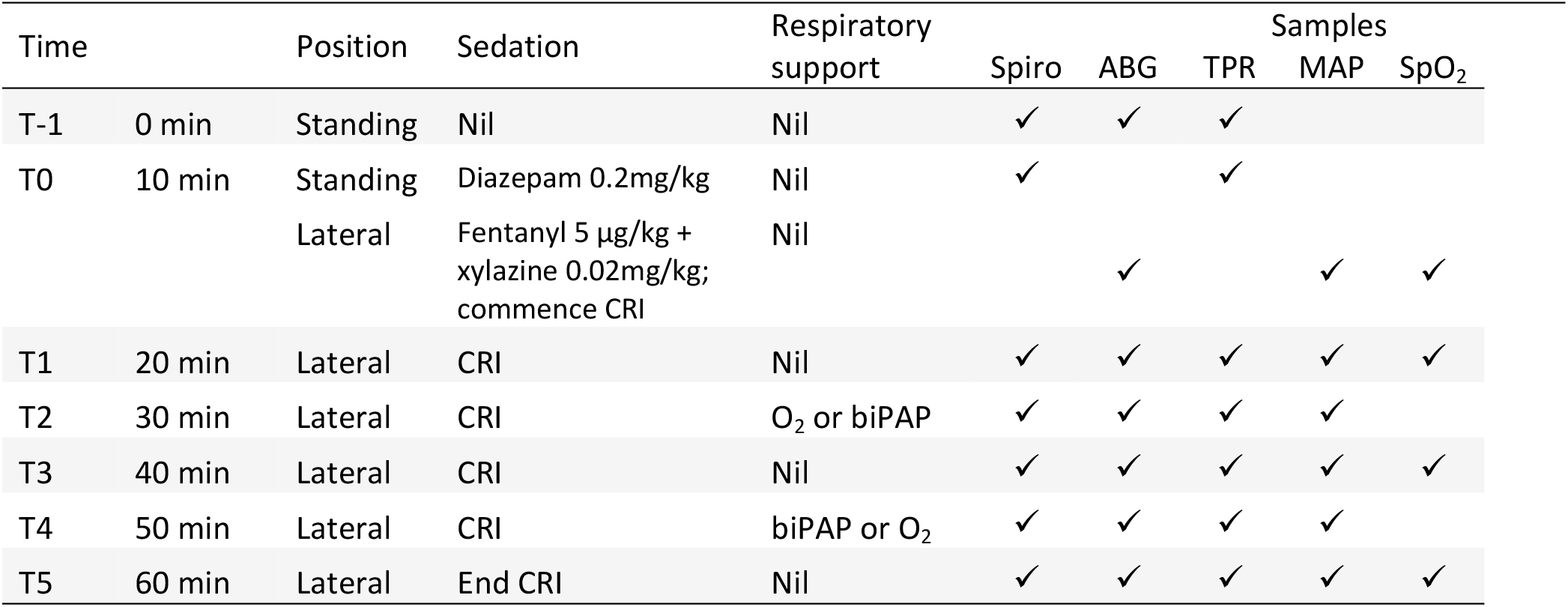
Treatment and sampling schedule for Group 3 foals. Baseline data were collected from standing, unsedated foals at T-1. Spirometry was performed at T0 in standing foals 5 minutes following administration of diazepam. Arterial blood gas, heart rate, respiratory rate and temperature were collected from recumbent foals within 5 to 10 minutes of the administration of a bolus injection of fentanyl and xylazine. A continuous infusion of fentanyl and xylazine in 0.9% sodium chloride delivered via syringe pump was commenced at the end of T0 (10 minutes lateral recumbency). Foals were randomised, in pairs, to receive oxygen administration (8L/min, O_2_) or non-invasive ventilation (biPAP at inspiratory pressure of 15 cmH_2_O and expiratory pressure at 5 cmH_2_O, with oxygen administration at 8L/min) at T2, with the reciprocal treatment administered at T4. Treatment order (biPAP / oxygen administration) was reversed in the second replicate. Spiro = spirometry and inspired/expired gas analysis; ABG = arterial blood gas: TPR = temperature, heart (pulse) rate, respiratory rate and mean arterial pressure; SpO_2_, pulse oximetry.

| Time |  | Position | Sedation | Respiratory support | Spiro | ABG | TPR | Samples MAP | SpO<sub>2</sub> |
| --- | --- | --- | --- | --- | --- | --- | --- | --- | --- |
| T-1 | 0 min | Standing | Nil | Nil | ✓ | ✓ | ✓ |  |  |
| T0 | 10 min | Standing | Diazepam 0.2mg/kg | Nil | ✓ |  | ✓ |  |  |
|  |  | Lateral | Fentanyl 5 µg/kg + xylazine 0.02mg/kg; commence CRI | Nil |  | ✓ |  | ✓ | ✓ |
| T1 | 20 min | Lateral | CRI | Nil | ✓ | ✓ | ✓ | ✓ | ✓ |
| T2 | 30 min | Lateral | CRI | O<sub>2</sub> or biPAP | ✓ | ✓ | ✓ | ✓ |  |
| T3 | 40 min | Lateral | CRI | Nil | ✓ | ✓ | ✓ | ✓ | ✓ |
| T4 | 50 min | Lateral | CRI | biPAP or O<sub>2</sub> | ✓ | ✓ | ✓ | ✓ |  |
| T5 | 60 min | Lateral | End CRI | Nil | ✓ | ✓ | ✓ | ✓ | ✓ |

Respiratory support (biPAP or O_2_ supplementation) was delivered via the large veterinary anaesthesia mask used for spirometry measurements. The mask was connected to a vented non-rebreathing elbow valve (Oracle 2 Vented Non-rebreathing Valve, 400HC206, Fisher and Paykel Healthcare, Nunawading, Victoria, Australia), and hence to a commercial, bi-level pressure support ventilator, as described for Group 1 foals. Oxygen delivery (8 L/min) was inserted into the system between the non-rebreathing valve and the ventilator tubing, as shown (Supplementary Figure S1). A pressure manometer (Advanced Anaesthetic Services, Gladesville, Australia) was connected to the biPAP ventilator to enable circuit pressure monitoring. Spirometry and gas sampling were performed by inserting the respiratory flow head and gas sampling port between the one-way valve and mask, as shown in Supplementary Figure 1, at the end of each respiratory intervention and after collection of arterial blood samples. Temperature, HR, RR and MAP were recorded over the final two minutes of each respiratory intervention. Pulse oximetry (SpO_2_) could not be performed during mask administration of O_2_ or biPAP because the transmission probe could not be placed on the tongue.

### Statistical methods

Power analysis from previous studies demonstrated that a sample size of six foals would discriminate differences in PaO_2_ and PaCO_2_ of 15 mmHg and 5mmgHg, respectively, with a power > 0.80 and α=0.05. Results from Group 1 and Group 2 foals are presented as raw data only due to the small number of foals in each group. A cross-over design was selected for the interventional study to further increase statistical power and to control for individual differences and possible effects attributable to treatment order. Data were tested for normality by the Shapiro-Wilks test and explored using appropriate descriptive statistics. The effect of replicate (Phase 1 or Phase 2) and sequence (O_2_ or biPAP at T2 with reciprocal treatment at T4) were evaluated by fitting separate mixed effects models using restricted maximum likelihood (REML) with time and replicate or sequence as random factors and foal as a fixed factor. In the absence of significant replicate or sequence effects, treatment effects (biPAP vs O_2_) were determined by mixed effects models with time as a random factor and subject as a fixed factor and post-hoc testing by Tukey’s method. Non-parametric results were analysed by Kruskal-Wallis test, with post-hoc testing by Dunn’s method. Relationships between PaO_2_, sO_2_ and pulse oximetry (SpO_2_), and between maximum CO_2_ in expired air (CO_2_max) and PaCO_2_, were explored using Pearson’s correlation; Bland-Altman analyses were used to assess agreement between these indices. Unless specifically stated, data satisfied criteria for normality and parametric tests were used. Significance was accepted as P<0.05 and all analyses were performed using Graph Pad Prism 8.4.3 for Windows, GraphPad Software, San Diego, California USA, www.graphpad.com).

## Results

### Group 1 foals

Three foals (F2, F4, F5) displayed a beneficial response to initial biPAP treatment, evidenced by increased PaO_2_ and decreased PaCO_2_ (Table 1). One foal (F1) demonstrated an improved PaO_2_, but minimal change to PaCO_2_ associated with treatment. On second and third treatment, F2 demonstrated minimal response, whereas repeat treatment of F5 demonstrated a similar beneficial effect to that observed after the first treatment. Importantly, however, F3 demonstrated reduced PaO_2_ and increased PaCO_2_ following 10 minutes of biPAP treatment. These changes were rapidly reversed when biPAP was discontinued and the foal allowed to breathe room air. Changes in inspired gases did not appear to predict this foal’s response to treatment, but respiratory rate decreased from 28 bpm to 10 bpm and protracted periods of apnoea were noted during biPAP administration.

### Group 2 foals

Two foals (F8 and F2) demonstrated improved oxygenation following biPAP, but all had increased PaCO_2_ (Table 1). Minimal effect was seen for F6 on both the first and second occasion when biPAP was administered (without supplementary oxygen). A dramatic reduction in PaO_2_ and increased PaCO_2_ were evidenced by F7 after biPAP, but changes were rapidly corrected when the foal was allowed to breathe room air. Increased CO_2_ was evident in analysis of expired gases during mask O_2_ supplementation and biPAP, suggesting that some degree of rebreathing occurred for this foal. Respiratory and heart rates did not vary during mask O_2_ administration or biPAP for this foal, but MAP increased (from 99 to 108 mmHg during other interventions, to 134 mmHg during biPAP) and SpO_2_ decreased to 89% (from 94 to 97% at FiO_2_ 21%) during biPAP.

### Group 3 foals

No effects attributable to replicate were observed for any parameter. Effects associated with sequence (biPAP or supplementary O_2_ at T2) are shown in supplementary materials. Heart and respiratory rates were highest in unsedated foals, and temperature decreased significantly throughout the study period (from 38.5°C to 37.8°C). Mean arterial pressure did not change associated with time or treatment (Supplementary Figure S2).

Oxygenation (PaO_2_) was greater in unsedated foals (T-1) than observed in sedated foals at T0 (P=0.002) or T1 (P=0.004, Figure 1). The administration of supplementary oxygen alone or with biPAP was associated with significantly increased PaO_2_ in comparison to results at all other sampling times (all P<0.001), and results following biPAP were significantly greater than after O_2_ administration (P=0.021). Arterial CO_2_ (PaCO_2_) results were not normally distributed and were resistant to transformation. Values were lowest in unsedated foals at T-1, and differences at this time and at T0 were significant when compared to results following administration of supplementary O_2_ and following biPAP (all P<0.001, Figure 1); differences at other times were not significant. Results following supplementary O_2_ administration were not significantly different to those obtained after biPAP (P=1.000). Hypercapnia (PaCO_2_ > 60 mHg) was observed for two foals (F8 and F9) following O_2_ administration (60mmHg and 66 mmHg, respectively), and following biPAP on both occasions for F9 (60 mmHg and 68 mmHg). Changes to blood pH mirrored changes to PaCO_2_ and effects on lactate and blood glucose treatment attributable to treatment were not observed (Supplementary Figure 3). Blood glucose concentrations increased across all sampling times (likely due to administration of xylazine), with results at T3 (P=0.013) and T5 (P=0.008) significantly higher than at T0, as were results during O_2_ (P=0.008) and biPAP (P=0.016, data not shown).

**Figure 1:**
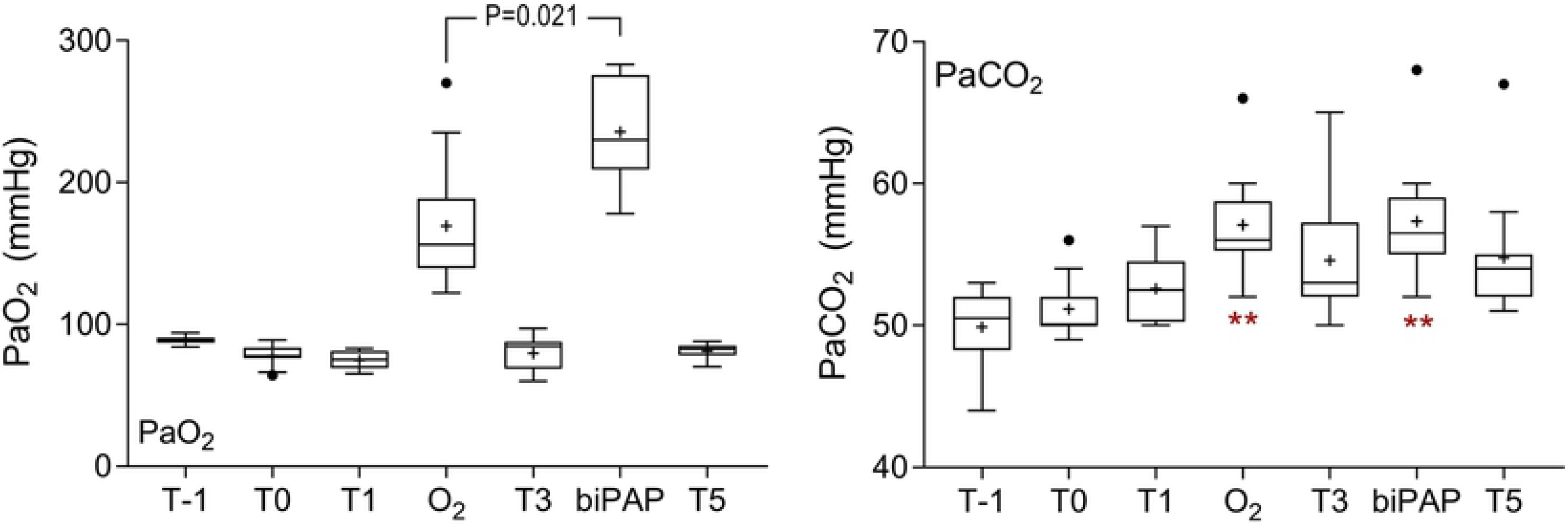
Blood gas results for Group 3 foals. Sedation was associated with a significant decrease in PaO_2_ at T0 (P=0.002) and T1 (P=0.004). The administration of supplementary oxygen by mask (O_2_) or during bi-level positive airway pressure ventilation (biPAP) was associated with a significant increase in PaO_2_ at all other time points (all P<0.001), and results following biPAP were significantly greater than following O_2_, as indicated. Results for arterial CO_2_ pressures were not normally distributed, and were resistant to transformation. Results at T-1 and T0 were significantly less than results following O_2_ and biPAP, as shown (**, P<0.001), following analysis by Kruskal Wallis test. Differences in PaCO_2_ were not different following O_2_ or biPAP (P=1.000). Data are shown as mean (+), median (horizontal line) and quartiles (box), with whiskers and outliers determined by Tukey method.

Significant time-sequence interactions were observed for spirometry variables including respiratory rate during spirometry (RRs), tidal volume (Vt), inspiratory time (Ti), expiratory time (Te) and peak inspiratory flow (PIF) (Supplementary Figure S4); biPAP was associated with significantly lower RR (P=0.014) and significantly longer inspiratory (P<0.001) and expiratory (P=0.020) times at T2 than observed following O_2_ administration at this time. Differences at other time points were not significant, so data have been combined for analysis of treatment effects. Sedation with diazepam at T0 was associated with a significant decrease in RRs relative to each other sampling times (Figure 2), except following biPAP (all P<0.050). Respiratory rate (RRs) during biPAP was lower than was observed at T1 (P=0.041), T3 (P=0.012) or following O_2_ administration (P<0.001). Tidal volume (Vt) was greatest in standing foals following diazepam sedation (T0) and values observed at this time and in unsedated foals at T-1 were significantly greater than observed in recumbent foals with the exception of during biPAP (all P<0.05). The administration of biPAP was associated with significantly greater Vt than was observed at T1 (P=0.001), T3 (P<0.001), T5 (P=0.011) and during O_2_ administration (P<0.001), and effects were more pronounced at T2 than at T4 (Supplementary Figure 4). Recumbency was associated with a significant reduction in minute ventilation relative to results from standing, unsedated foals (T-1), but this effect was not observed during biPAP (Figure 2).

**Figure 2:**
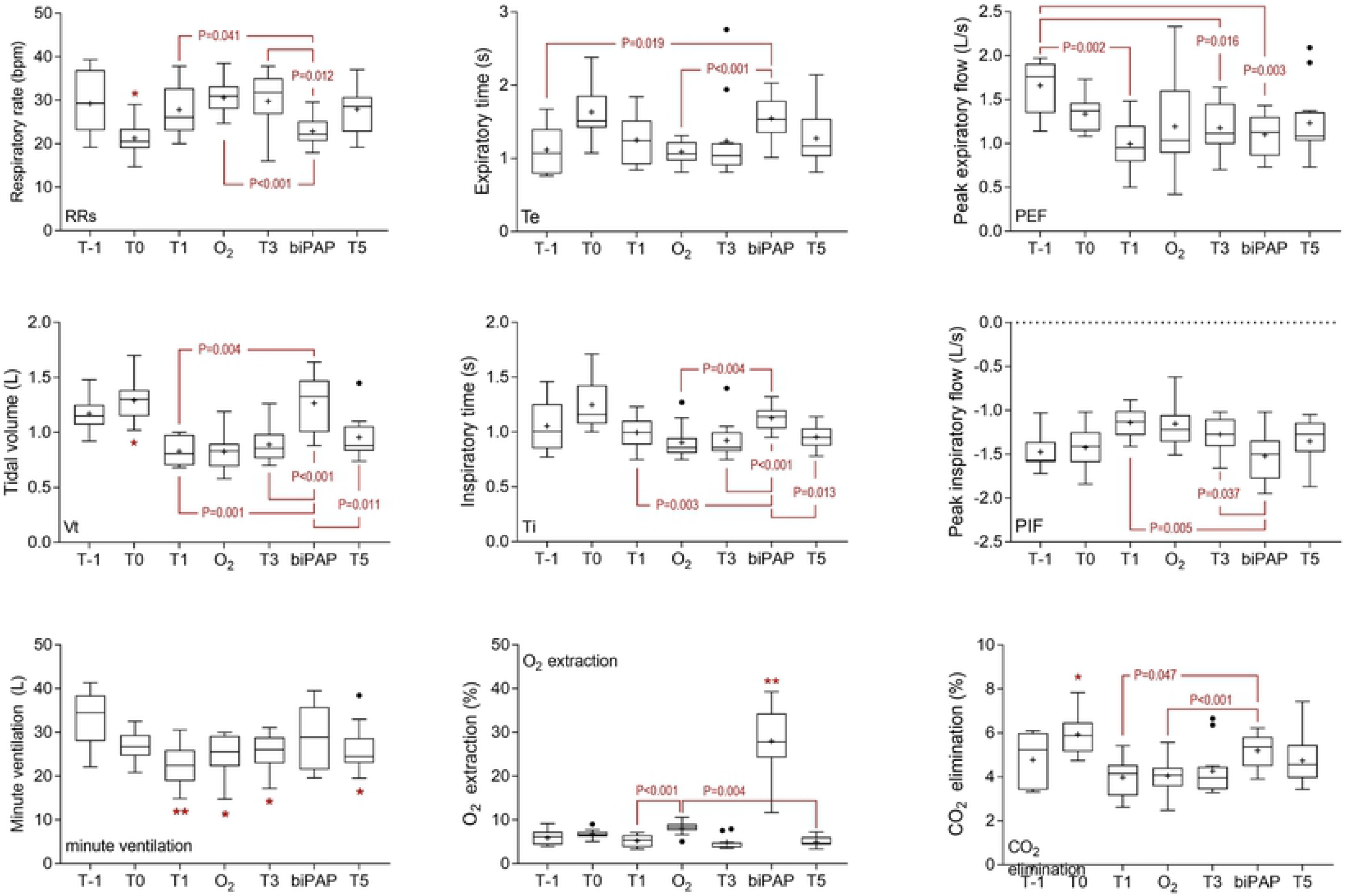
Spirometry and gas exchange results for Group 3 foals. Sedation was associated with a significant decrease in respiratory rate during spirometry (RRs) at T0, and this effect was significant (P<0.05) in comparison with results at all other time point except during bi-level positive airway pressure ventilation (biPAP). Effects on tidal volume (Vt) reciprocated those observed on RRs, with differences again observed at T0 (P<0.05 when compared to other sampling points with the exception of during biPAP). Minute ventilation was greatest in standing, unsedated foals (T-1), and significant decreases were observed at all other sampling points (*, P<0.05, **, P<0.01) except during biPAP. Inspiratory (Ti) and expiratory (Te) times were longest in standing foals following sedation with diazepam (T0), but significant effects were observed only in comparison with T5 (P=0.020) for Ti. For Te, comparisons between T0 and T-1 (P=0.008), O_2_ (P=0.006) and T5 (P=0.012) were significant. Significant time effects on peak expiratory (PEF) and inspiratory (PIF) flows are shown. The administration of biPAP was associated with greater O_2_ extraction than observed at any other time (**, all P<0.01). Oxygen extraction was also greater during mask O_2_ administration, as shown, and at T0 (standing foals following administration of diazepam) than at T3 (P=0.046) or T5 (P<0.001). The elimination of CO_2_ was greatest at T0 than at any other time, except following biPAP (*, all P<0.05). Differences between effects observed following biPAP administration and at other times are shown. Data are shown as mean (+), median (horizontal line) and quartiles (box), with whiskers and outliers determined by Tukey method.

Significant effects were observed for both inspiratory and expiratory time, reflective of changes observed in RRs (Figure 2), but there was no effect on I:E ratio (data not shown). Peak inspiratory flow was greatest during biPAP (−1.52 L/s), and significant effects were observed compared to values obtained at T1 (P=0.005) and T3 (P=0.037). Expiratory flows were greatest in unsedated foals (T-1), and significant differences were observed at T1 (P=0.002), T3 (P=0.016) and during biPAP (P=0.003).

Time and sequence effects were observed for data derived from analysis of inspired / expired gas composition due to differences following O_2_ administration or biPAP at T2 (Supplementary Figure S5). Differences at other time points were not significant, so data have been combined for analysis of treatment effects. As expected, the administration of supplementary O_2_ was associated with an increased FiO_2_ during both mask supplementation and biPAP, compared to FiO_2_ when breathing room air (all P<0.001), and values were significantly greater during biPAP than during administration of O_2_ only (P=0.004). Oxygen concentrations in expired air were also greater following the administration of supplementary O_2_, but differences were not observed between mask O_2_ administration and biPAP (P=0.172). Oxygen extraction was much greater during biPAP than at all other time points (all P<0.005), including during O_2_ administration (P<0.001, Figure 2). Oxygen extraction was also greater during mask O_2_ administration than at T1 (P<0.001) or T5 (P=0.004), and in standing foals at T0 relative to T3 (P=0.046) and T5 (P<0.001). Maximum concentrations of CO_2_ (CO_2_max) were observed at T0 in standing foals following administration of diazepam and observed differences were significant in comparison with results at T-1 (P=0.016), T1 (P=0.0003), T3 (P=0.026), T5 (P=0.0002) and following O_2_ administration (P=0.031). Results after biPAP were significantly greater than following O_2_ administration (P=0.047). Minimum concentrations of CO_2_ were observed during biPAP, and differences were significant in comparison with T3 (P=0.021) and T5 (P=0.013). The elimination of CO_2_ was greatest at T0 in diazepam sedated foals, with significant differences observed between results at T0 and all other sampling points, with the exception of biPAP (all P<0.030, Figure 2). Results observed following biPAP were significantly greater than observed at T1 (P=0.047) or following O_2_ administration (P=0.004).

### Monitoring

Paired results for pulse oximetry (SpO_2_) and haemoglobin saturation determined by blood gas analysis (sO_2_) were available for 47 data sets from the current study. SpO_2_ results correlated significantly with sO_2_ (r=0.61, 95% CI 0.34 to 0.78, P<0.001), but there was poor agreement between these two methods of assessing haemoglobin saturation (Figure 3). Although bias was minimal (1.4%, standard deviation 5.4%), the observed limits of agreement were large (−11.95 to 9.2%), and increased divergence was observed for results obtained from the most hypoxic foal. Paired results for CO_2_max and PaCO_2_ were available for 136 data sets (Figure 3). There was poor but significant correlation between results (r=0.25, 95% CI 0.09 to 0.40, P=0.003), and agreement was poor (bias 14.0 ± 8.9%) with broad limits of agreement (−3.5 to 31.5%).

**Figure 3:**
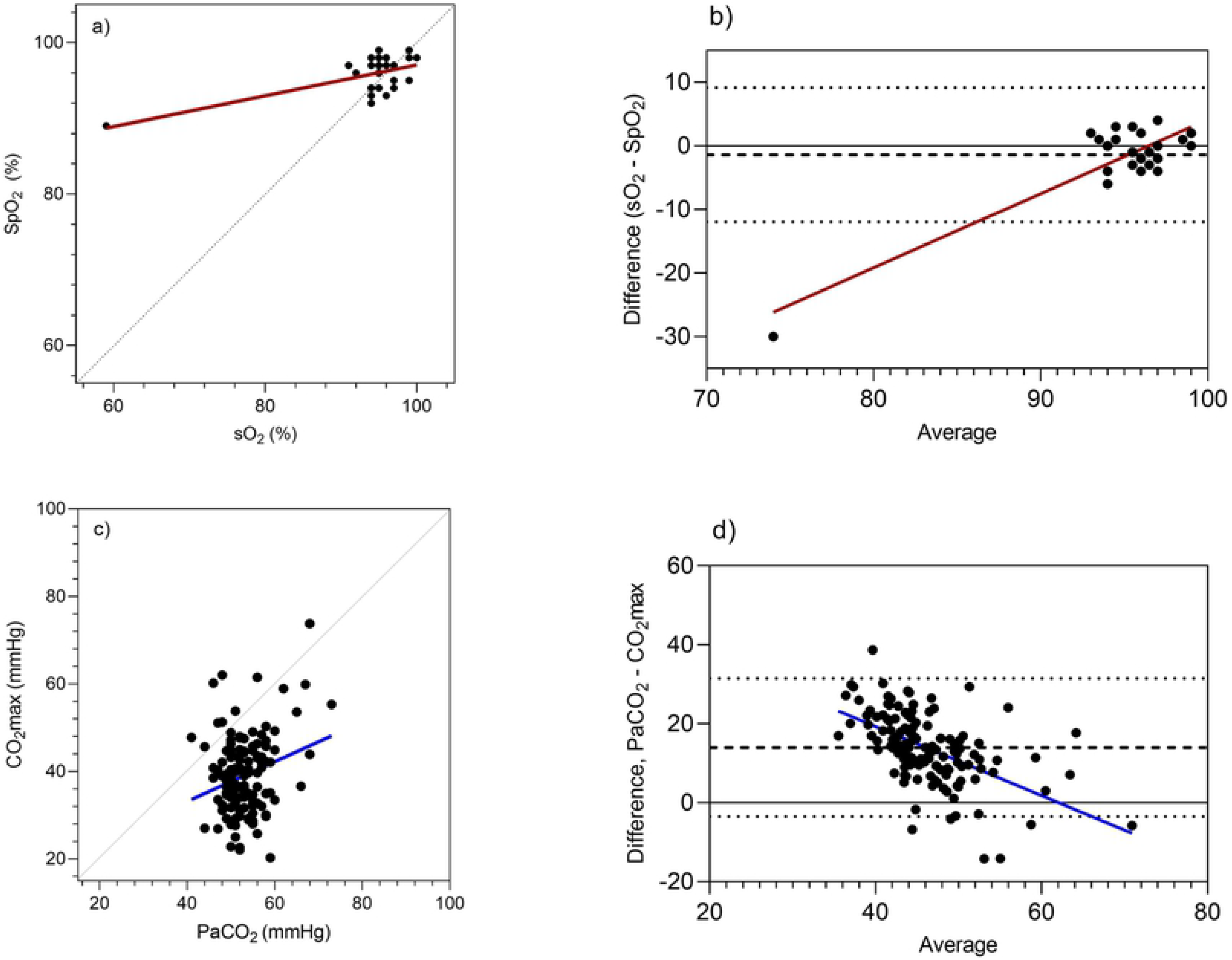
Associations between blood gas results and non-invasive measures of oxygenation (pulse oximetry, SpO_2_) and carbon dioxide accumulation (CO_2_max). Results are presented as correlations (a and c), with perfect agreement (unity) shown as a dotted line. Agreement is shown following Bland-Altman analysis (b and d), with the mean difference (dashed line) and limits of agreement (dotted lines) shown. Oxygenation of haemoglobin (sO_2_) and partial pressure of CO2 (PaCO_2_) were determined from blood gas analyses.

## Discussion

The administration of biPAP in the current study was associated with a positive response to treatment in four of five foals with respiratory disease (Group 1), and two of four healthy, sedated neonatal foals < 28 days age (Group 2). However, one healthy foal (F6) showed minimal response to initial treatment, and one foal in Group 1 showed minimal response to subsequent treatments (F2’, F2’’). More importantly, one foal with respiratory insufficiency (F3, Group 1) and one healthy Group 2 foal (F7) demonstrated hypoxia and hypercapnia following biPAP treatment. For F3 this was associated with decreased respiratory rate, suggesting inadequate alveolar ventilation might have caused this outcome. Conversely, for F7 this was associated with increased expired CO_2_ during both mask O_2_ administration and biPAP, suggesting that rebreathing or other mechanisms for CO_2_ accumulation within equipment dead space might have contributed to hypercapnia in this case. Of clinical parameters that might have been observed in the absence of CO_2_ monitoring, only MAP was increased for this foal.

Results from Group 3 foals demonstrated that improved blood oxygenation was achieved following both mask administration of supplementary O_2_ and biPAP. Benefits associated with biPAP were greater, and were also associated with decreased RR, increased Vt and increased minute ventilation. Increased Vt represents a more efficient ventilation strategy than increased RR, as there is increased alveolar ventilation relative to ventilation of airway dead space, and decreased RR is likely to be associated with decreased work of breathing. Increased inspiratory pressure during biPAP was associated with increased PIF, and the adverse effects on PEF observed during CPAP in previous studies (13) were not observed in the current study, presumably due to the lower expiratory pressures during biPAP ventilation. As expected, increased FiO_2_ was observed during mask administration of supplementary O_2_ and during biPAP, so observed increases in arterial oxygenation likely reflect the steeper diffusion gradient resulting from these changes. Surprisingly, FiO_2_ was higher during biPAP than during O_2_ administration to Group 3 foals. This was not observed for Group 1 or 2 foals, and is not expected during positive pressure treatments where increased flow is associated with decreased partial pressure of O_2_ (27). As the minimum inspired CO_2_ was greater during the administration of supplementary O_2_ than during biPAP, it is likely that mask administration of supplementary O_2_ in the current study was associated with CO_2_ retention, a problem avoided by the use of a non-rebreathing valve during biPAP, and by nasal insufflation of oxygen for Group 1 foals.

Increased PaCO_2_ was observed in Group 3 foals following respiratory support, including hypercapnia (PaCO_2_ > 60 mmHg) following O_2_ administration (two foals) or biPAP (one foal). Hypercapnia has been reported in response to O_2_ supplementation in human neonates (28) and foals (13, 29), and may be due to reduced respiratory drive, increased metabolic rate, hypoventilation due to sedation or effects of equipment dead space. As noted above, mask administration of supplementary O_2_ was associated with the accumulation of CO_2_ within equipment dead space (rebreathing) in the current study, but this was not observed during biPAP. Despite decreased RR, minute ventilation during biPAP was the same as observed in standing, unsedated foals in the current study suggesting that biPAP prevented reduced ventilation associated with sedation and recumbency. However, our hypothesis, that lower expiratory pressures and improved expiratory function would ameliorate hypercapnia was not demonstrated. As was observed during CPAP (13), the observed increases in PaCO_2_ and pH in the current study were modest, and consistent with current ventilation strategies that accept increased arterial CO_2_ tension and hypercapnic acidosis (‘permissive hypercapnia’) as acceptable consequences without adverse effects on outcome (30, 31), and with possible therapeutic effects (32). Sedation, and the supraphysiological PaO_2_ values observed in the current study, might have contributed to the observed hypercapnia in Group 2 and Group 3 foals. With the exception of one foal with markedly reduced RR, hypercapnia was not observed in Group 1 foals, where sedation was not required for respiratory support. Alternatively, Group 1 foals, with spontaneous respiratory insufficiency, might have been less susceptible to adverse effects of ventilatory support. A similar difference has been noted in human patients with recruitable air spaces (indicated by the presence of a lower inflection point in the inspiratory pressure-volume (PV) curve), in comparison to patients with no such inflection (21). Although we were unable to monitor PV curves in our study, it is possible that Group 1 foals with spontaneous disease responded more appropriately to the imposition of PEEP than Group 2 and 3 foals with healthy lungs where the increased expiratory pressure might be more likely to cause pulmonary overdistension.

Although reversed within 10 minutes of cessation of biPAP, the adverse effects of biPAP on F3 (Group 1) and F7 (Group 2), and the hypercapnia observed in Group 3 foals, demonstrate the necessity for close monitoring during the implementation of respiratory support in equine neonates. Evaluation of NIV should consider effects on both oxygenation and carbon dioxide. Direct measurement of PaO_2_ and sO_2_ by co-oximetry is more accurate than blood gas analysis for determination of oxygenation (33), but neither technique provides an immediate result or allows for continuous monitoring. Arterial samples can be difficult to obtain in hypovolaemic foals or animals with distal limb oedema, and risks associated with arterial puncture include pain, haemorrhage, arterial injury, aneurism formation, thrombosis and distal ischaemia. Pulse oximetry has been recommended as an appropriate alternative to invasive sampling for determination of haemoglobin saturation (34). Previous studies have suggested that placement of transmission or reflectance sensors on the lip or tongue ensures the most reliable assessment of SaO_2_ in foals (34), although bias and limits of agreement in that study were similar to those observed in the current study and well outside accepted standards of care (35). Probe placement on the tongue was not possible during mask administration of respiratory support in the current study, and was not tolerated by unsedated foals.

End-tidal CO_2_ (PETCO_2_) is commonly monitored during anaesthesia as an indirect measure of PaCO_2_. Whilst PETCO_2_ has been reported as an acceptable technique for monitoring of neonatal foals (34), studies during NIV in people have suggested that, as observed in the current study, the technique was not predictive of PaCO_2_ or changes in PaCO_2_ (36). The association between PETCO_2_ and PaCO_2_ assumes that the patient exhales fully, and that end-expiratory gases approximate gas composition in the alveoli. This assumption is not valid if, as we have hypothesised, our ventilated foals are not exhaling completely. For this reason peak expired CO_2_ concentrations have been termed FeCO_2_max in the current study. Alternative techniques to assess ventilatory function, such as volume capnography and electrical impedance tomography (37), offer greater capacity to more accurately assess response to treatment.

A number of limitations are noted in the current study. The number of foals presented for management of respiratory disease was low and, for three of five foals, the degree of respiratory insufficiency was mild. Findings in healthy foals with pharmacologically-induced respiratory insufficiency may not be predictive of responses in neonates with spontaneous disease. We were unable to document alveolar ventilation, physiological dead space or PEEP*i* in the current study, which was designed to evaluate a readily available intervention and monitoring strategies that might be readily implemented in a practice setting. The assessment of non-invasive monitoring was based on a small number of observations, and the data set did not include results from foals with severe hypoxaemia or hypercapnia.

## Conclusions

Consistent with previous studies evaluating CPAP, biPAP was an effective respiratory support strategy for healthy foals with pharmacologically induced respiratory insufficiency and for a small number of foals with spontaneous respiratory disease. BiPAP was associated with increased PaO_2_, more efficient gas exchange and changes in respiratory mechanics including increased tidal volume, decreased respiratory rate and increased peak inspiratory flow. The technique preserved minute ventilation in the face of reductions associated with sedation and recumbency observed at other times, but was associated with modest increases in PaCO_2_. As in previous studies, the use of a commercially available ventilator intended for at-home care of adults with chronic obstructive respiratory conditions or sleep apnoea represents an available and potentially cost effective option for use in equine practice, although there is a need for careful and frequent monitoring of patient oxygenation and ventilation during NIV. Our results suggest that monitoring of alveolar ventilation, PV curves and PEEP*i* might be important for effective NIV of foals and to better characterise the response of foals to respiratory support. The use of lower expiratory pressures in the current study did not prevent hypercapnia, and increases in PaCO_2_ observed in the current study were similar to those observed during CPAP in healthy foals in previous studies. Similar increases were not observed in the majority of foals with respiratory disease suggesting that foals with pulmonary pathology might respond differently to foals with pharmacologically or centrally-induced respiratory suppression. Observed effects on were PaCO_2_ were rapidly reversed and predominantly within acceptable bounds for permissive hypercapnia. Although not a primary objective of the current study, our results suggest the non-invasive monitoring approaches used in this study were not reliable, and techniques are needed for more accurate, non-invasive assessment of respiratory function in foals during NIV.

## Acknowledgements

This study was funded by AgriFutures Australia. Numerous veterinary and equine science students assisted with restraint and care of foals (and their dams) including Cathrine Borgen-Nielsen, Tegan Davis, Philippa Kellett and Stacey Walker. Special thanks go to Equine Centre staff at CSU for their dedicated and ongoing care of mares and foals, and to Jaymie Loy and Equine Science reproduction students for their care of mares and foals at foaling. ResMed Pty Ltd donated bi-level positive airway (biPAP) ventilators used in this study, and Jeff Armistead provided technical advice. This project was completed in partial fulfilment of Veterinary Honours for Lexi Burgemeestre.

